# ADAR1 editing dependency in triple-negative breast cancer

**DOI:** 10.1101/2020.01.31.928911

**Authors:** Che-Pei Kung, Kyle A. Cottrell, Sua Ryu, Emily R. Bramel, Raleigh D. Kladney, Emily A. Bross, Leonard Maggi, Jason D. Weber

**Author notes:** These authors contributed equally (Che-Pei Kung, Kyle A. Cottrell).

## Abstract

Triple-negative breast cancer (TNBC) is the deadliest form of breast cancer. Unlike other types of breast cancer that can be effectively treated by targeted therapies, no such targeted therapy exists for all TNBC patients. The ADAR1 enzyme carries out A-to-I editing of RNA to prevent sensing of cellular double-stranded RNAs (dsRNA). ADAR1 is highly expressed in breast cancer including TNBC. Here, we demonstrate that ADAR1 expression and editing activity is required in TNBC cell lines but not in ER+ and/or Her2+ cells. In TNBC cells, knockdown of ADAR1 attenuates proliferation and tumorigenesis. PKR expression is elevated in TNBC and its activity is induced upon ADAR1-knockdown, which correlates with a decrease in translation. ADAR1-dependent TNBC cell lines also exhibit elevated IFN stimulated gene expression. IFNAR1 reduction significantly rescues the proliferative defects of ADAR1 loss. These findings establish ADAR1 as a novel therapeutic target for TNBC tumors.

## Introduction

Generally defined by the lack of estrogen receptor (ER), progesterone receptor (PR) and HER2 expression, triple-negative breast cancer (TNBC) accounts for 15 to 20 percent of all breast cancer diagnoses in the United States each year(Ademuyiwa et al., 2017). Unlike ER-positive (tamoxifen, fulvestrant, and other ER modulators) and HER2-positive (Herceptin and other HER2 inhibitors) breast cancers, there are no targeted therapies for all TNBC patients(Waks and Winer, 2019). The lack of targeted therapies for TNBC leaves chemotherapy as the main treatment option that carries a generally worse prognosis(Garrido-Castro et al., 2019). Efforts to develop effective targeted therapies against TNBC have focused on further sub-categorizing TNBC based on gene expression signatures, as well as looking to exploit common genetic vulnerabilities(Perou, 2011, Anders et al., 2016).

A potential therapeutic target for TNBC is Adenosine Deaminase Acting on RNA (ADAR1, encoded by *ADAR*). ADAR1 caries out the enzymatic reaction of deaminating adenosine to inosine within cellular dsRNA, in a process known as A-to-I editing. Induction of ADAR1 expression is prevalent in breast cancer(Fumagalli et al., 2015, Han et al., 2015, Paz-Yaacov et al., 2015, Peng et al., 2018, Anantharaman et al., 2017) and ADAR1-mediated A-to-I editing has been found to influence the levels of its targets in breast cancer(Gumireddy et al., 2016, Binothman et al., 2017, Dave et al., 2017, Nakano et al., 2017). Recent studies indicate that ADAR1 is over-represented in TNBC and may be correlated with poor prognosis when RNA editing is increased(Song et al., 2017, Sagredo et al., 2018).

ADAR1 acts in a negative feedback loop to inhibit the type-I IFN pathway triggered by endogenous dsRNAs or dsRNAs introduced upon viral infections(Mannion et al., 2014, Liddicoat et al., 2015). ADAR1 has been shown to suppress type-I IFN pathway through multiple mechanisms, including destabilization of the dsRNA structure, reduced expression, and activation of the dsRNA sensors MDA5 and RIG-I, and inhibition of IFN expression(Mannion et al., 2014, Liddicoat et al., 2015, Pestal et al., 2015, George et al., 2016, Li et al., 2012, Pujantell et al., 2017). ADAR1-mediated A-to-I RNA editing by the IFN-inducible p150 isoform (not the constitutive p110 isoform) is essential for its ability to modulate dsRNA-induced IFN signaling(Liddicoat et al., 2015, Pestal et al., 2015, George et al., 2016). ADAR1’s ability to regulate this response was recently linked to the development of ADAR1 dependency in some cancer cell lines; two groups showed that by removing ADAR1 from cancer cells with elevated innate immune signaling, cells became susceptible to inflammation-induced cell death(Gannon et al., 2018, Liu et al., 2019). This is consistent with previous findings that ADAR1 prevents immune and translational catastrophes by blocking dsRNA-activated IFN pathway(Mannion et al., 2014, Chung et al., 2018).

Here we demonstrate that TNBC cell lines are dependent on ADAR1 expression and activity; loss of ADAR1 in these cell lines inhibits cellular growth and tumorigenesis, highlighting the therapeutic potential of ADAR1 inhibitors for the treatment of TNBC.

## Results

### ADAR1 is highly expressed in all breast cancer subtypes

Using publicly available data from TCGA (The Cancer Genome Atlas)(Han et al., 2015, Fumagalli et al., 2015), we found that high expression of ADAR1 correlated with poor prognosis of breast cancers (Figure 1A). Recent studies indicated that ADAR1 promotes tumorigenesis of metaplastic breast cancers, and that high expression of ADAR1 correlates with poor prognosis in basal-like breast cancers(Sagredo et al., 2018, Dave et al., 2017). Since both basal-like and metaplastic breast cancers share similar characteristics with TNBC, we sought to determine the importance of ADAR1 in the tumorigenesis of TNBC. By evaluating the TCGA database, we found that while mRNA expression of ADAR1 was higher in TNBC compared to normal, it was not significantly different between TNBC and non-TNBC tumors (Figure 1B). Additionally, ADAR1 expression was not significantly higher in any one subtype of breast cancer based on PAM50 classification(Lehmann et al., 2016) (Supplemental Figure 1A). This observation is consistent with data from the Cancer Cell Line Encyclopedia (CCLE), which uses both RNA-seq and Reverse Phase Protein Array (RPPA) to determine RNA and protein expression levels in numerous cancer cell lines (Supplemental Figure 1B-C). Data from both the TCGA and CCLE datasets also revealed that both p150 and p110 isoforms of ADAR1 were expressed at similar levels between TNBC and non-TNBC specimen (Supplemental Figure 1D-H), with p110 expression being consistently higher than p150 in all samples. Additionally, we assessed p150 isoform expression by immunohistochemistry in TNBC and non-TNBC patient tumors, Figure 1D. We sought to determine the protein expression level of the ADAR1-p150 isoform in a panel of established breast cancer cell lines representing TNBC and non-TNBC. Immunoblot analysis showed that ADAR1 (p150 isoform) is overexpressed, compared to normal human mammary epithelial cells (HMECs), in over half of all TNBC (6/8) and non-TNBC (5/8) cell lines assayed (Supplemental Figure 1I-K). These results indicate that ADAR1-p150 is overexpressed in many breast cancer cell lines regardless of subtype.

**Figure 1:**
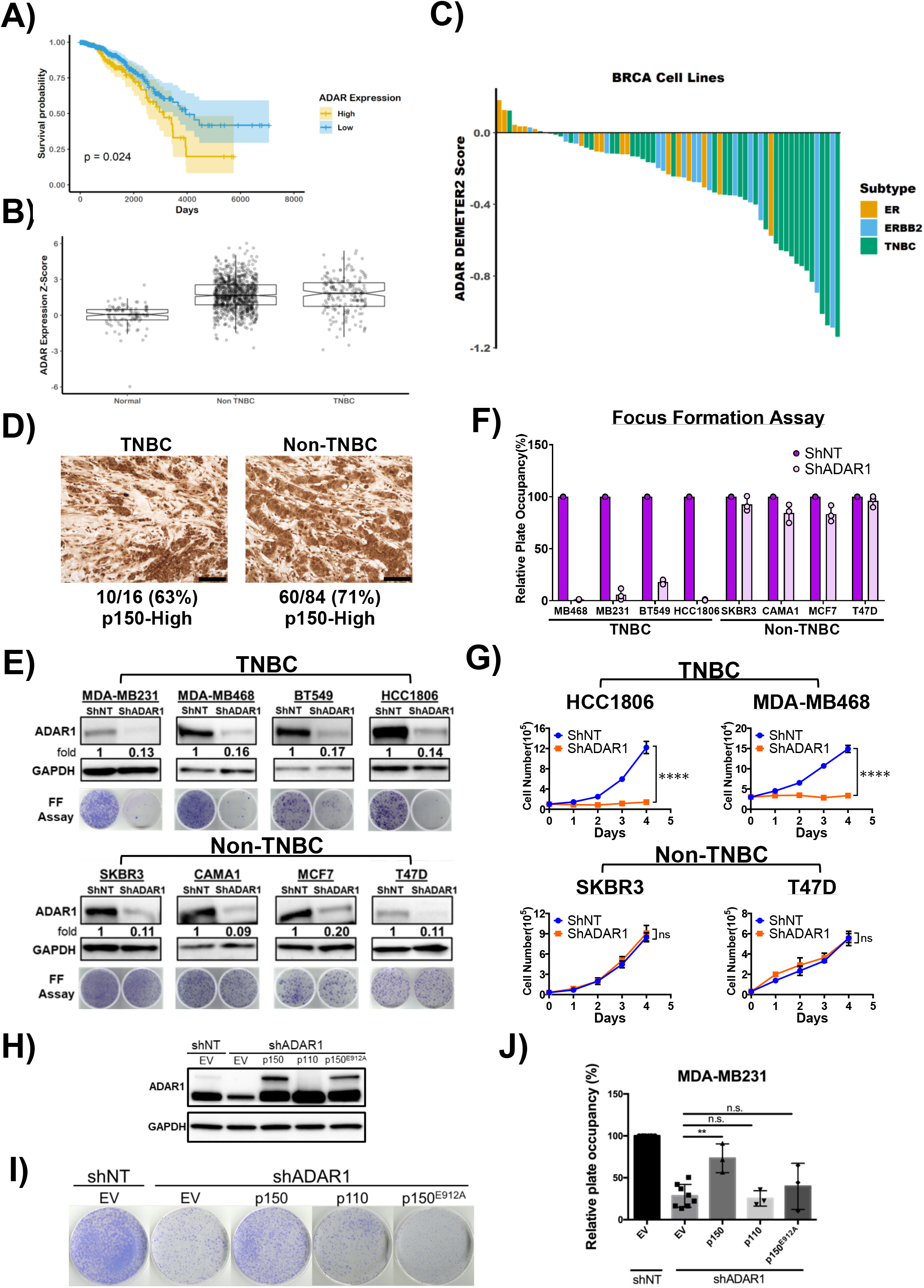
ADAR1 is highly expressed in all breast cancer subtypes and required for TNBC proliferation. **A)** Kaplan-Meier survival curves of breast cancer patients. Patients were stratified by ADAR1 expression, above or below z-score = 2.34. **B)** Relative mRNA expression of ADAR1 in normal, TNBC and Non-TNBC breast cancer. Data were extracted from TCGA database. **C)** ADAR1-dependency scores in breast cancer cell lines. Lower DEMETER2 scores indicate stronger ADAR1-dependency. ERBB2 = HER2-positive. **D)** Representative images of IHC staining of ADAR1 in TNBC and Non-TNBC breast cancer tissues (scale: 100 μM). Numbers below the image indicate the ratio of samples identified as high p150-ADAR based on IHC scoring. **E)** Immunoblots showing protein levels of ADAR1 p150 isoform and GAPDH(loading control) with or without ADAR1-knockdown in breast cancer cell lines. Fold change of ADAR1 (ShADAR1/ShNT) is indicated, normalized to GAPDH. Focus formation (FF) assay showed that ADAR1-knockdown reduced proliferation of TNBC but not Non-TNBC cells. Images are representative, N=3. **F)** Quantification of FF in **E)**. Relative plate occupancy was determined using ImageJ software and normalized to ShNT samples for each cell line. Data are represented as mean ± SD, N=3. **G)** Cell proliferation assay showing that ADAR1-knockdown reduced proliferation of TNBC but not Non-TNBC cells. Data are represented as mean ± SD, N=2. (****) p<0.0001. ns, not significant. **H)** Immunoblots showing protein levels of ADAR1 and GAPDH(loading control) with overexpression of p150, p110 or editing-defective p150^E912A^ in ShADAR1-treated MDA-MB231 cells. Images are representative, N=3. EV, empty virus. **I)** FF assay showing that p150, but not p110 or editing-defective p150^E912A^, partially rescued proliferation of ShADAR1-treated MDA-MB231. Images are representative, N=3. **J)** Quantification of FF in **I)**. Relative plate occupancy was determined using ImageJ software and normalized to ShNT-EV. Data are represented as mean ± SD, N>3. (**) p<0.01. n.s., not significant. See also Figure S1 and S2.

### ADAR1 is required for TNBC proliferation

Several recent studies have suggested that some established cancer cell lines display strong dependencies on ADAR1 expression(Liu et al., 2019, Ishizuka et al., 2019, Gannon et al., 2018). Given the high expression of ADAR1-p150 in most breast cancer cell lines, we sought to determine whether these breast cancer cell lines exhibited ADAR1-dependency. We analyzed publicly available RNAi and CRISPR-Cas9 datasets to determine if ADAR1 was required for the survival of breast cancer cell lines representing various subtypes(McFarland et al., 2018, Meyers et al., 2017). TNBC and basal-like cell lines made up the majority of breast cancer cells exhibiting high ADAR1 sensitivity scores (DEMETER2 Score < −0.5) (Figure 1C, Supplemental Figure 2A-C). Importantly, we did not observe a correlation between ADAR1 expression and ADAR1-dependency across these breast cancer cell lines (Supplementary Figure 2D). To experimentally validate ADAR1-dependency among breast cancer cell lines, we knocked-down ADAR1 expression in eight cell lines (Four TNBC: MDA-MB231, MDA-MB468, BT549, HCC1806; Four non-TNBC: SKBR3, CAMA1, MCF7, T47D); all of these cell lines showed noticeable ADAR1-p150 isoform overexpression over HMEC controls in our immunoblot analysis (Supplemental Figure 1I). Long-term (7-28 days) and short-term (4 days) cell proliferation was evaluated for each cell line following ADAR1 knockdown. Notably, similar levels of ADAR1 knockdown were achieved for each cell line (Figure 1E). All four TNBC cell lines displayed significant attenuation in both long- and short-term proliferation following ADAR1 knockdown (Figure 1F-G, Supplemental Figure 2E). Conversely, ADAR1 expression proved dispensable for proliferation in all four non-TNBC cell lines.

### ADAR1-p150 editing activity rescues TNBC proliferation

While both isoforms of ADAR1 are expressed in TNBC, our knockdown experiment does not distinguish between p150 or p110 dependence. To address this, we set up a knockdown-rescue system. We overexpressed either the p110 or p150 isoform following ADAR1 knockdown in MDA-MB231 cells and evaluated their ability to rescue cell proliferation (Figure 1H-J). Overexpression of ADAR1-p150, but not p110, resulted in significant rescue of cell proliferation in MDA-MB231 TNBC cells. Having established a rescue system for ADAR1 dependent proliferation, we next aimed to determine whether the editing activity of ADAR1-p150 was required for this rescue. An editing-defective mutant (E912A) of the p150 isoform was incapable of rescuing the ADAR1 knockdown phenotype, indicating that the A-to-I editing function of ADAR1 is absolutely required for cellular proliferation in TNBC cells (Figure 1I-J).

### ADAR1 is required for TNBC transformation and tumorigenesis

To assess the functional relevance of our findings, we investigated the requirement of ADAR1 for the transformation of breast cancer cell lines. We utilized anchorage independent growth in soft agar as a measure of cellular transformation. Knockdown of ADAR1 dramatically reduced soft agar colonies of MDA-MB231 and HCC1806 TNBC cells while not significantly affecting the numbers of colonies formed by SKBR3 and T47D non-TNBC cells (Figure 2A-D).

**Figure 2:**
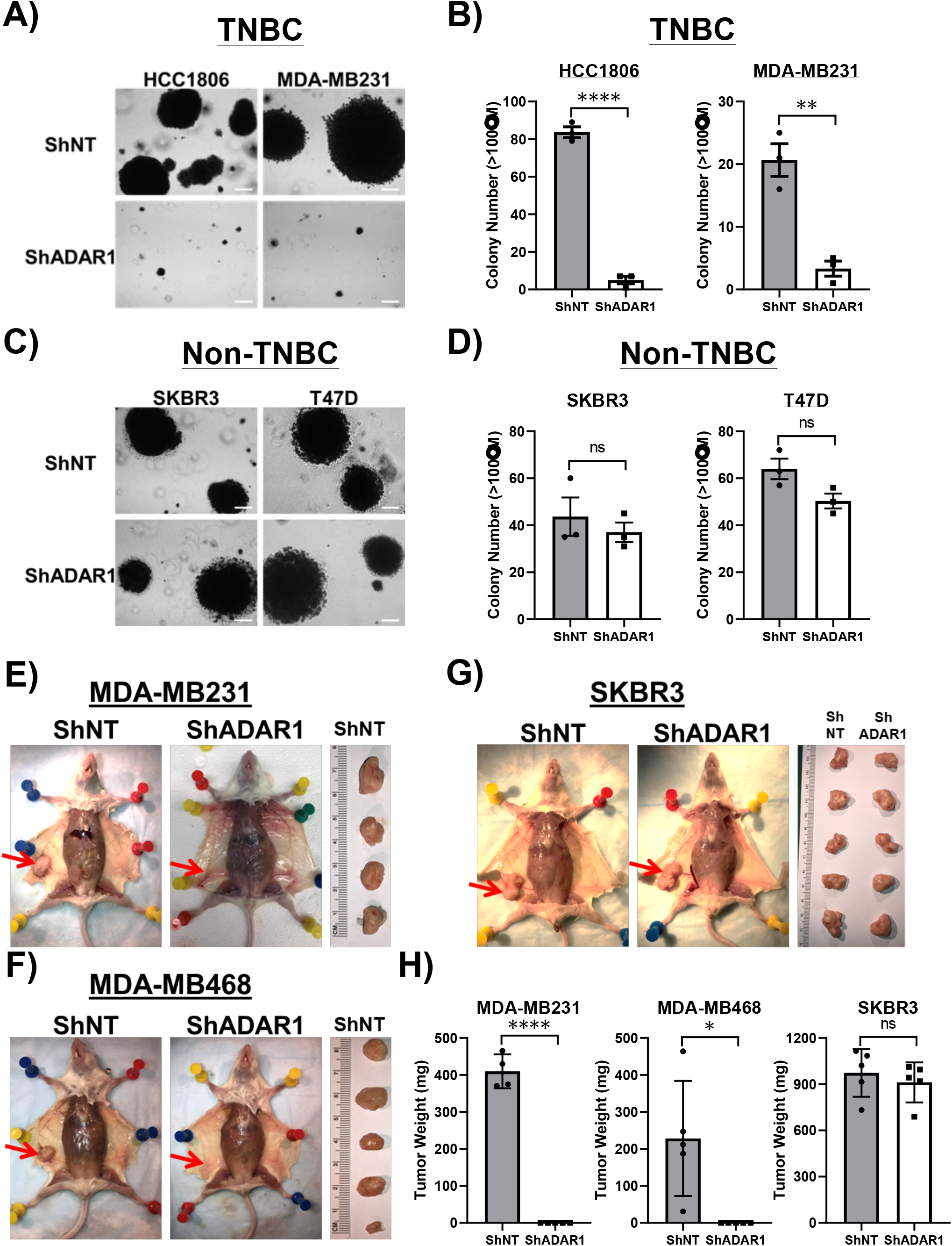
ADAR1 is required for TNBC transformation and tumorigenesis. **A)** Soft agar assay (SAA) showing that ADAR1-knockdown reduced anchorage-independent growth of TNBC cells (HCC1806 and MDA-MB231). Images are representative, N=3. Scale-bar, 100μM. **B)** Quantification of SAA in **A)**. Colonies bigger than 100μM in diameter were counted. Data are represented as mean ± SD, N=3. (****) p<0.0001. (**) p<0.01. **C)** SAA showing that ADAR1-knockdown did not affect anchorage-independent growth of Non-TNBC cells (SKBR3 and T47D). Images are representative, N=3. Scale-bar, 100μM. **D)** Quantification of SAA in **C)**. Colonies bigger than 100μM in diameter were counted. Data are represented as mean ± SD, N=3. ns, not significant. **E)** Orthotopic implantation of MDA-MB231 cells into abdominal mammary fat pad. Tumors were removed from the mice ~4 weeks post injection and weighed (ShNT, N=4; ShADAR1, N=5). Red arrows indicate the location of mammary fat pad. **F)** Orthotopic implantation of MDA-MB468 cells into abdominal mammary fat pad. Tumors were removed from the mice ~12 weeks post injection and weighed (N=5). Red arrows indicate the location of mammary fat pad. **G)** Orthotopic implantation of SKBR3 cells into abdominal mammary fat pad. Tumors were removed from the mice ~4 weeks post injection and weighed (N=5). Red arrows indicate the location of mammary fat pad. **H)** Quantification of the result shown in **E)–G)**. Data are represented as mean ± SD. (****) p<0.0001. (*) p<0.05. ns, not significant.

To extend these *in vitro* findings, we next determined whether ADAR1 was required for TNBC cell lines to form tumors *in vivo*. We performed mammary gland orthotopic transplantations using TNBC and non-TNBC cells following ADAR1 knockdown. Parental (shRNA-NonTargeting, shNT) MDA-MB231 and MDA-MB468 TNBC cells and SKBR3 non-TNBC cells were all able to form visible tumors in the mammary glands of four-five independently transplanted female immune compromised mice (Figure 2E-H). Knockdown of ADAR1 in MDA-MB231 and MDA-MB468 TNBC cells completely abrogated their ability to form tumors in transplanted mice. In contrast, ADAR1 knockdown in SKBR3 cells did not significantly affect tumor formation in transplanted mammary glands. Collectively, these results demonstrate that ADAR1 expression is required for *in vitro* transformation and *in vivo* tumor formation of TNBC cells, but is completely dispensable for these properties in non-TNBC cells.

### PKR is overexpressed in TNBC and activated upon ADAR loss

Previous reports have shown that ADAR1 dependency in human cancer cells could be mediated through several downstream pathways, including translational inhibition triggered by activated PKR and ribonuclease L (RNASEL), as well as type-I IFN signaling(Li et al., 2017, Gannon et al., 2018, Liu et al., 2019). To investigate if these pathways contribute to the ADAR1 dependency observed in TNBC cells, we first analyzed the TCGA and CCLE datasets to determine if these pathways are intrinsically elevated in TNBC. Across TCGA breast cancer samples, RNA expression of PKR is significantly higher in TNBC samples compared to non-TNBC (Figure 3A). This is consistent with RNA-seq data for breast cancer cell lines within the CCLE (Figure 3B). Moreover, elevated PKR expression positively correlates with the ADAR1 sensitivity scores, suggesting a strong relationship between PKR and TNBC-associated ADAR1 dependency (Figure 3C, Supplemental Figure 3A-C). We further confirmed this observation by immunoblot analysis among our panel of sixteen breast cancer cell lines to show a general elevation of PKR expression across all TNBC cell lines (Figure 3D). We also detected heightened levels of PKR phosphorylation as well as its downstream substrate eIF2α in TNBC cells compared to non-TNBC cells. Upon ADAR1 knockdown, phosphorylation of PKR and eIF2α was markedly induced in all TNBC cell lines but remained unchanged in the non-TNBC cell lines (Figure 3E). These observations suggest that TNBC-associated ADAR1 dependency might be attributed to PKR-mediated translational inhibition. To investigate this, we performed polysome profiling. ADAR1 knockdown in MDA-MB231 and HCC1806 TNBC cells led to inhibition of translation, demonstrated by the substantial reduction of polysome peaks (Figure 3F-G). These results suggest that translational repression contribute to TNBC-associated ADAR1 dependency. While attempting to rescue the ADAR-knockdown phenotype in MDA-MB231 and HCC1806 cells by knockdown of PKR, we observed that knockdown of PKR alone greatly reduced foci formation (Supplemental Figure 3D-E). This suggests basal PKR expression is required for the proliferation of these cell lines and precluding us from determining if expression of PKR is required for the ADAR-knockdown phenotype.

**Figure 3:**
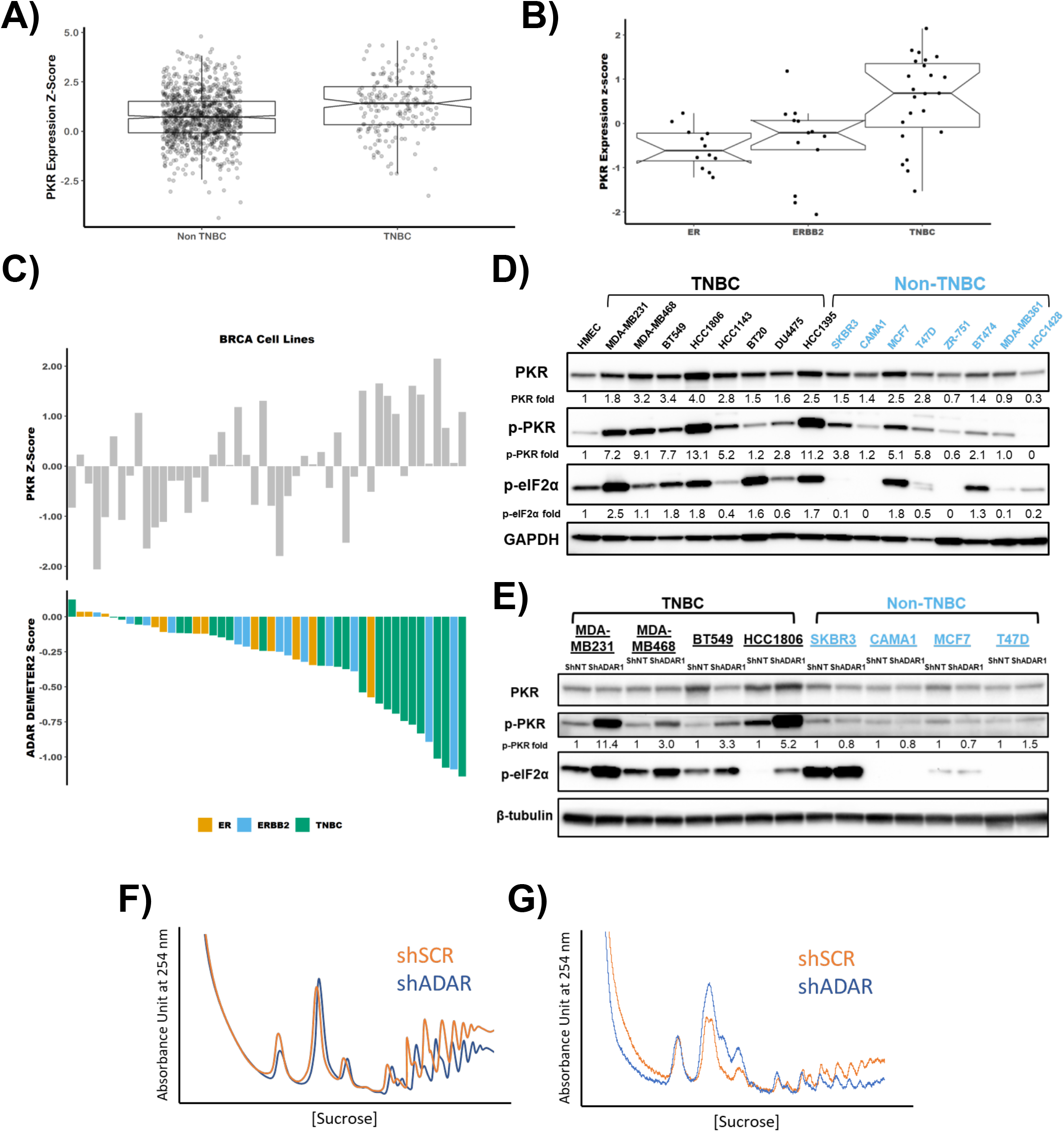
PKR is overexpressed in TNBC and activated upon ADAR loss. **A)** Relative mRNA expression of PKR in TNBC and Non-TNBC. Data were extracted from TCGA database. **B)** Relative mRNA expression of PKR in ER-positive, ERBB2(HER2)-positive and TNBC cell lines. Data were extracted from CCLE database. **C)** ADAR1-dependency scores positively correlate with PKR expression. Upper panel: PKR expression z-score in breast cancer cell lines. Lower panel: ADAR1-dependency scores. Lower DEMETER2 scores indicate stronger ADAR1-dependency. **D)** Immunoblots showing protein levels of PKR, p-PKR (T446), p-eIF2α (S51) and GAPDH(loading control) in breast cancer cell lines. Densitometry quantification of gel images was normalized to GAPDH and set relative to HMEC signal. Data shown are representative, N=3. **E)** Immunoblots showing protein levels of PKR, p-PKR (T446), p-eIF2α (S51) and β-tubulin (loading control) in TNBC and non-TNBC breast cancer cell lines with or without ADAR1-knockdown. Densitometry quantification of gel images was normalized to GAPDH and compared to HMEC signal set as 1-fold. Data shown are representative, N=3. **F)** Polysome profiling of MDA-MB231 cells with or without ADAR1-knockdown. Data shown are representative of three replicates. **G)** Ribosomal profiling of HCC1806 cells with or without ADAR1-knockdown. See also Figure S3.

### RNASEL is not activated following loss of ADAR1 in TNBC

Activation of RNASEL and subsequent translational inhibition has also been shown to result in cell lethality in the absence of ADAR1(Li et al., 2017). The CCLE dataset indicated that RNASEL activators OAS1, OAS2 and OAS3 were highly expressed in ADAR1 dependent cell lines, while the expression of *RNASEL* showed modest correlation with ADAR1 dependency (Supplemental Figure 3F-G). A hallmark of RNASEL activation is degradation of rRNA(Silverman et al., 1983). However, we did not observe rRNA degradation in ADAR1-dependent TNBC cells after ADAR1 knockdown (Supplemental Figure 3H), further suggesting that the RNASEL pathway does not significantly contribute to TNBC-associated ADAR1 dependency and the induction of OAS genes likely reflects the fact that OAS genes are also known ISGs (see below).

### ADAR1-dependent TNBCs exhibit elevated ISG expression

Another factor contributing to ADAR1 dependency in cancer cell lines is the type-I IFN pathway(Liu et al., 2019). It has been shown previously that this connection is mediated through either altering the expression of type I IFN regulators or activating the feed-forward loop of IFN signaling(Gannon et al., 2018, Liu et al., 2019). RNA expression data from the TCGA and CCLE datasets showed that TNBC have higher ISG expression (Core ISG Score(Liu et al., 2019)) compared to non-TNBC (Figure 4A-B). This is consistent with the elevated expression of PKR and ISG15 in our immunoblot analysis among breast cancer cell lines (Figure 3D, Supplemental Figure 4A). Like PKR expression, the Core ISG Score positively correlated with ADAR1 sensitivity among TNBC cell lines (Figure 4C and Supplemental Figure 4B-C).

**Figure 4:**
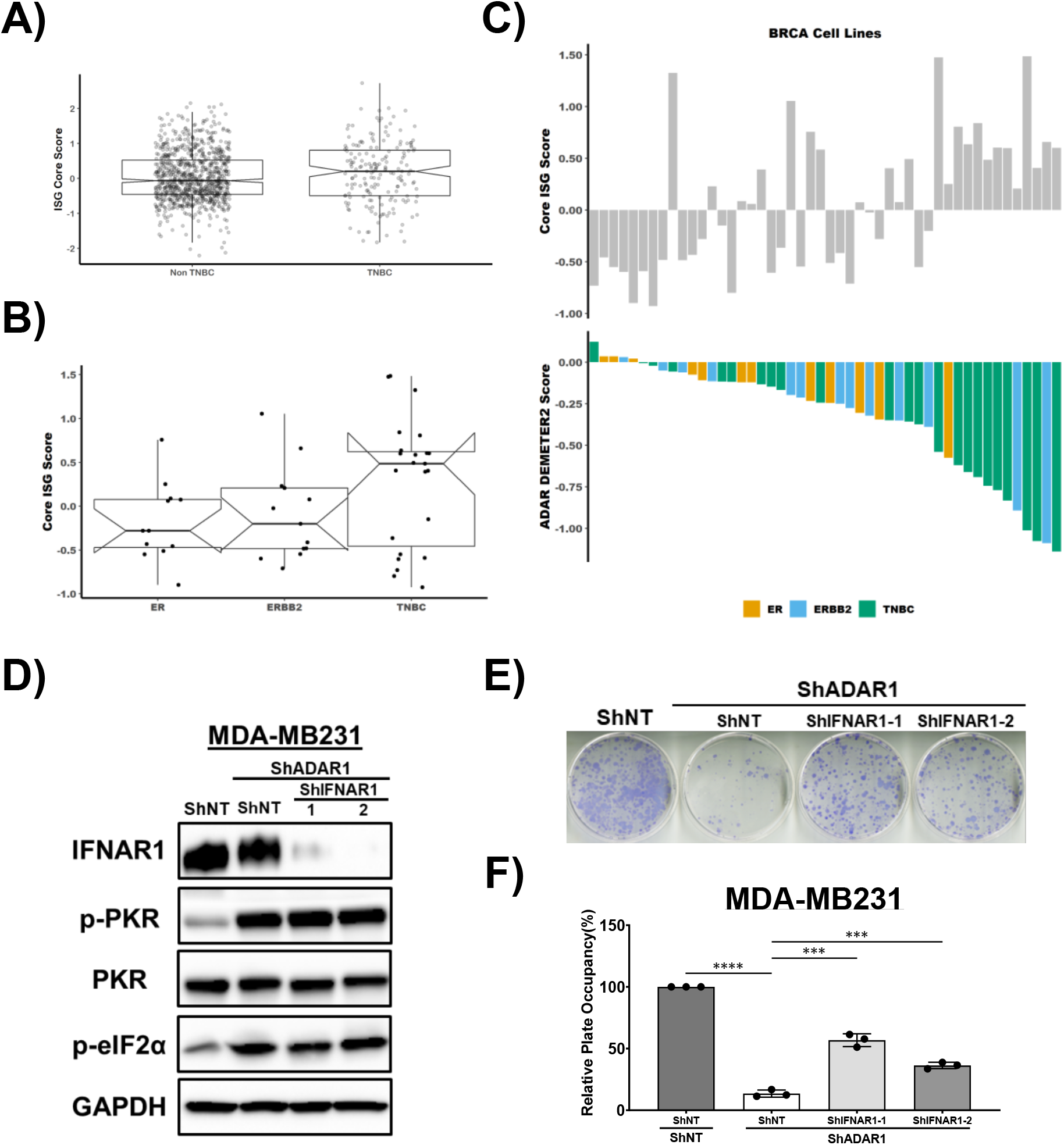
ADAR1-dependent TNBCs exhibit elevated ISG expression and INFAR1 loss rescues ADAR1 knockdown phenotype. **A)** Relative ISG Core Scores in TNBC and Non-TNBC breast cancer samples. Data were extracted from TCGA database. **B)** Relative ISG Core Scores in ER-positive, ERBB2(HER2)-positive and TNBC cell lines. Data were extracted from CCLE database. **C)** ADAR1-dependency scores positively correlate with ISG Core Scores in breast cancer cell lines. Upper panel: ISG Core Scores in breast cancer cell lines. Lower panel: ADAR1-dependency scores. Lower DEMETER2 scores indicate stronger ADAR1-dependency. **D)** Immunoblots showing protein levels of IFNAR1, PKR, p-PKR (T446), p-eIF2α (S51) and GAPDH(loading control) in MDA-MB231 cells. IFNAR1 was knocked down in ShADAR1-treated MDA-MB231 cells to determine if IFNAR1 loss reverses ADAR1-knockdown phenotype. Images are representative, N=3. **F)** FF assay showing that IFNAR1 loss partially rescued ADAR1-knockdown phenotype in MDA-MB231 cells. Images are representative, N=3. **G)** Quantification of FF in **F)**. Relative plate occupancy was determined using ImageJ software and normalized to ShNT-ShNT. Data are represented as mean ± SD. N=3. (****) p<0.0001. (***) p<0.001. See also Figure S4.

### INFAR1 loss rescues ADAR1 knockdown phenotype

To establish whether the type-I IFN pathway accounts for the significant differences of ADAR1-dependency between TNBC and non-TNBC cell lines, non-TNBC cell lines (SKBR3 and MCF7) were treated with IFNβ in ADAR1-intact and ADAR1-deficient cells (Supplemental Figure 4D-G). Expression of ADAR1 and ISG15 were induced upon IFNβ treatment, as well as phosphorylation of STAT1. However, while the treatment of IFNβ generally reduced cell proliferation, it did not sensitize non-TNBC cells to ADAR1 deficiency (Supplemental Figure 4E and G), implying that IFNβ alone is not capable of switching ADAR1-resistant cells to ADAR1-dependent cells.

To determine if the type-I IFN pathway functionally contributes to ADAR1 dependency in TNBC, we knocked-down ADAR1 and the IFN alpha-receptor subunit 1 (IFNAR1) simultaneously in both MDA-MB231 and MDA-MB468 cells (Figure 4D and Supplemental Figure 4H). The knockdown of IFNAR1 partially rescued the proliferation of both cell lines, suggesting that TNBC-associated ADAR1 dependency can be partially attributed to type I IFN pathway (Figure 4E-F, Supplemental Figure 4I). However, knockdown of IFNAR1 in TNBC cells did not alter the levels of phosphorylated PKR (Figure 4D, Supplemental Figure 4H), suggesting that in these TNBC cells, either type-I IFN and PKR pathways independently contribute to ADAR1 dependency or IFNAR1 resides downstream of PKR.

## Discussion

Recent studies have highlighted the dependence of some cancer cell lines on ADAR1 expression(Gannon et al., 2018, Liu et al., 2019). Here, we characterized the requirement for ADAR1 in a panel of established breast cancer cell lines. ADAR1-dependent cell lines shared an elevated ISG-expression signature. Loss of ADAR1 in these cell lines led to activation of the translational regulator PKR and translational repression. The ADAR1-dependence phenotype could be partially abrogated by knockdown of IFNAR1. It is not currently understood what makes select cancer cell lines ADAR1-dependent, or conversely why others are refractory to ADAR1-loss. It has been proposed that the higher ISG expression might potentiate these cells towards ADAR1-dependency – loss of ADAR1 would further elevate ISG expression leading to the growth inhibition phenotype(Liu et al., 2019, Gannon et al., 2018). However, we have demonstrated that for cell lines refractory to ADAR1 loss, treatment with IFN-β did not render them sensitive to ADAR1 knockdown. Furthermore, we observed no activation of PKR in the ADAR1 refractory cell lines following ADAR1 loss. These findings suggest that the link between ADAR1 loss and the IFN pathway or PKR activation in ADAR1-refractory cell lines is missing. Loss of ADAR1 is thought to activate the IFN pathway and PKR by causing an increase in dsRNA – stemming from a reduction in A-to-I editing(Mannion et al., 2014, Liddicoat et al., 2015). It is possible that ADAR1-refractory cell lines either do not accumulate dsRNA following ADAR1 loss or there exists a system that prevents dsRNAs from activating the IFN pathway or PKR. Understanding the molecular basis of this process would help to predict which cell lines – or more importantly which tumors – should be sensitive to ADAR1 loss.

Important clinical implications can be drawn from these observations. Our data suggest that ADAR1 is a legitimate candidate for targeted therapies in TNBC. We found that TNBC cell lines and patient samples exhibit elevated ISG and PKR expression, which is consistent with ADAR1-dependent cell lines. With increased understanding of ADAR1 functions, novel therapeutic strategies against ADAR1 could benefit ADAR1-dependent cancers, including TNBC(Kung et al., 2018). Secondly, the relationship between ADAR1 dependency and type-I IFN pathway could point to new directions for TNBC interventions. Recent studies revealed that the increased IFNβ target gene signature correlates with improved recurrence-free survival in TNBC, and IFNβ treatment inhibits tumor progression in TNBC by reducing cancer stem cell (CSC) plasticity(Doherty et al., 2017, Doherty et al., 2019). In addition to cell-intrinsic effects of ADAR1-loss in cancer cells, removal of ADAR1 has been shown to sensitize tumors to immunotherapy by overcoming resistance to checkpoint blockade(Ishizuka et al., 2019).

It was recently demonstrated that chemotherapies elicit a state of immunological dormancy in ER-negative breast cancers, marked by sustained type-I IFN signaling, reduced cell growth, and longer progression-free survival(Lan et al., 2019). This indicates a possible shared mechanism between chemotherapy-induced immunological dormancy and ADAR1-dependency in TNBC. It is important to note that careful considerations need to be given when applying the concepts of ADAR1 inhibition and type-I IFN application in the treatment of TNBC. It is recognized that type-I IFN can elicit paradoxical effects on cancer development(Snell et al., 2017). For example, it has been suggested that type-I IFN pathway, potentially through ISG15-mediated ISGylation, can promote the aggressiveness of TNBC(Forys et al., 2014, Lo et al., 2018). Therefore, further understanding of the relationship between ADAR1 functions and TNBC tumorigenesis should better inform the context in which this strategy can provide the maximum benefit.

## Acknowledgments

This work was supported by R01CA190986 (JDW), F32GM131514 (KAC) and TL1TR002344 (C-PK) from the National Institute of Health, and W81XWH-18-1-0025 from the Department of Defense (JDW). This work was supported by the Longer Life Foundation: A RGA/Washington University partnership. The results shown here are in whole or part based upon data generated by the TCGA Research Network: https://www.cancer.gov/tcga.

## Author Contributions

Conceptualization, C-PK, KAC, and JDW; Methodology, C-PK, KAC, and JDW; Software, KAC; Investigation, C-PK, KAC, SR, ERB, RK, EAB, LM, and JDW; Writing – Original Draft, C-PK and KAC; Writing – Review & Editing, C-PK, KAC, SR, ERB, RK, LM, and JDW; Funding Acquisition, JDW; Supervision, JDW

## Experimental Procedures

### Cell lines and reagents

Human mammary epithelial cells (HMEC) and breast cancer cells lines were obtained from American Tissue Cells Consortium (ATCC). HMECs were cultured in MammaryLife Basal Medium (Lifeline Cell Technology) and passaged by using 0.05% trypsin-EDTA (Gibco) and Defined Trypsin Inhibitor (DTI, Gibco). All breast cancer cell lines were maintained in Dulbecco’s Modification of Eagle’s Medium (DMEM, GE Life Sciences) supplemented with 10% fetal bovine serum (Gibco, 10091-148), Sodium Pyruvate (Cellgro, 30-002-CI), Non-Essential Amino Acids (NEA, Cellgro, 25-030-CI), and L-glutamine (Cellgro, 25-005-CI). Lipofectamine 2000 (Invitrogen) was used for transfection to generate lentivirus. Fugene 6 transfection reagent (Roche) was used for all other transfection experiments.

### Immunoblot analysis

Cell lysates were extracted from cells at ~90% confluence. Cell were washed with phosphate-buffered saline (PBS, GE Life Sciences), scrape harvested, centrifuged at 1000 × g for 5 min, and lysed with RIPA buffer (20mM Tris-HCl pH7.5, 150mM NaCl, 1mM EDTA, 1% NP-40, 0.1% SDS, 0.1% Deoxycholate) supplemented with 10mM PMSF and HALT protease inhibitor cocktail (Thermo Fisher Scientific). Lysates were clarified by centrifugation and the protein concentration was determined using DC protein assay system (Bio-Rad Laboratories). Equal amount of protein was resolved by sodium dodecyl sulfate-polyacrylamide gel electrophoresis (SDS-PAGE) using Criterion TGX Stain-Free Precast Gels (Bio-Rad) and transferred onto Immobilon-P membranes (MilliporeSigma). Primary antibodies used in this study include ADAR1 (Santa Cruz, sc-73408), MDA5 (Cell Signaling, #5321), RIG-I (Cell Signaling, #3743), PKR (Cell Signaling, #3072), PKR Thr-446-P (Abcam, ab32036), STING (Cell Signaling, #13647), IFNAR1 (Bethyl Laboratories, A304-290A), ISG15 (Santa Cruz, sc-166755), GAPDH (Bethyl Laboratories, A300-641A), β-Tubulin (Abcam, ab6046), EIF2S1/eIF2α Ser-51-P (Abcam, 32157), EIF2S1 (Abcam, ab5369). Secondary antibodies conjugated to Horseradish peroxidase were used at a dilution of 1:5-10,000 (Jackson Immunochemicals). Clarity Western ECL Substrate (Bio-Rad) was then applied to blots and protein levels were detected using autoradiography with ChemiDoc XRS+ Imager (Bio-Rad). Densitometry quantification of protein signals was performed using ImageJ software (NIH, Bethesda, MD).

### Quantitative reverse-transcription polymerase chain reaction

Total RNA was isolated from cells using RNeasy Mini Plus kit (Qiagen, Hilden, Germany) including on-column DNase digestion following the manufacturer’s protocol. High Capacity cDNA Reverse Transcription Kit (Life technologies, CA, USA) was used to transcribe RNA to cDNA. Quantitative PCR (qPCR) was performed using iTaq Universal SYBR Green Supermix (Bio-Rad, #1725121) on the C1000 Thermal Cycler (CFX96 Real-Time System, Bio-Rad), and data analysis was performed using the 2(-ΔΔC_T_) method. Messenger RNA expression levels were normalized to GAPDH. Primers used in this study are listed in Supplemental Table 1.

### Lentiviral production and transduction

To generate lentivirus, transformed human embryonic kidney HEK293T cells were transfected using Lipofectamine 2000 (Invitrogen) with pCMV-VSV-G, pCMV-ΔR8.2, and expression constructs (with pLKO.1-puromycin or pLKO.1-hygromycin backbone for short-hairpin RNAs and with pLVX-hygromycin backbone for overexpression constructs). Growth medium was replaced with fresh medium 24hr after transfection, and supernatants containing lentivirus were harvested 24hr later. For transduction, one million cells were infected with lentivirus for 24hr in the presence of 10 μg/ml protamine sulfate to facilitate viral entry. Sequences of shRNAs are listed in Supplemental Table 1. ADAR1 overexpression constructs (p150; p110; p150-E912A mutant) were subcloned from pBac-ADAR1 constructs (generous gifts from Kazuko Nishikura at The Wistar Institute, Philadelphia) into pLVX-hygromycin vectors (Cho et al., 2003).

### Cell proliferation and Focus Formation Assays

For cell proliferation assays, 2-5X10^4^ cells were plated in triplicate in six-well plates. Cells were trypsinized, harvested and counted using a hemocytometer or the Cello Cell counter every 24 hours for 4 days post-plating. For the focus formation assay, 3-5X10^3^ cells were plated in triplicate in 10-cm cell culture dishes 7-28 days (depending on the cell line: MDA-MB231 and BT549, ~7 days; HCC1806, ~14 days; SKBR3, ~14-21 days; T47D, ~21 days; MDA-MB468, CAMA1 and MCF7, ~28 days) post-plating until foci became visible. Cells were washed with PBS twice, fixed with 100% methanol, dried, and stained with Giemsa staining reagent (Sigma Aldrich). Stained plates were scanned and surface areas occupied by cell foci were measured using ImageJ software (NIH, Bethesda, MD).

### Soft agar transformation assay

Equal volumes of 2X concentrated DMEM culture media and 1% noble-agar solution (made with sterile cell-culture-grade water) were mixed to make 0.5% agar-media solution and plated in the bottom of six-well plates. Equal volumes of 2X concentrated DMEM culture media and 0.6% noble-agar solution were mixed to make 0.3% agar-media solution for cell suspension. 2-5X10^4^ cells were suspended in 0.3% agar-media solution and layered, in triplicate, onto the bottom layer. Cells were fed with fresh media every 7 days and incubated in 37°C for 21-30 days, before being stained with 0.005% crystal violet and examined under a microscope. Colonies bigger than 100 μM in diameter were manually counted.

### Mammary gland orthopedic implantation

The abilities of human breast cancer cell lines to form tumors *in vivo* were evaluated by performing mammary gland orthopedic implantation as described previously (Brenot et al., 2018). Immuno-deficient NOD scid gamma (NSG) female mice at 6-8 week-old were purchased from Jackson Laboratory (Bar Harbor, ME) and used for this experiment. 1-3X10^6^ cells were harvested and resuspended in PBS, mixed with standard base-membrane Matrigel Matrix (Corning, MA, USA) at 1:1 volume ratio, and kept at 4°C until implantation. In total 100μl of cells-Matrigel solutions were injected into the right inguinal mammary glands of NSG mice, which were monitored closely to observe tumor formation. Mice were euthanized before tumors in control groups reached 2 cubic cm in size, and palpable tumors were dissected from the mice for weight measurement. All animal-related experimental procedures were performed in compliance with the guidelines given by the American Association for Accreditation for Laboratory Animal Care and the U.S. Public Health Service Policy on Human Care and Use of Laboratory Animals. All animal studies were approved by the Washington University Institutional Animal Care and Use Committee (IACUC) in accordance with the Animal Welfare Act and NIH guidelines (Protocol 20160916)

### Statistical analysis

Unless otherwise stated, the two-tailed unpaired Student t test was performed for statistical analysis. All in vitro and in vivo data are reported as the mean ± SD unless stated otherwise, Statistical analyses were performed using GraphPad Prism. P values are as indicated: *, p < 0.05; **, p < 0.01; ***, p < 0.001; ****, p < 0.0001; n.s., not significant.

### Polysome profiling

Either MDA-MB231 or HCC1806 cells were treated with 100 μg/mL cycloheximide in growth media for 5 minutes at 37 °C. The cells were washed with ice-cold 1x PBS containing 100 μg/mL cycloheximide prior to harvesting by scraping. The cells were lysed in polysome lysis buffer (20 mM Tris pH 7.26, 130 mM KCl, 10 mM MgCl_2_, 0.5% NP-40, 0.2 mg/mL heparin, 200 U/mL RNasin, 2.5 mM DTT, 1x HALT, 100 μg/mL cycloheximide, 0.5% sodium deoxycholate) for 20 minutes on ice prior to clarification at 8000 g for 10 minutes at 4 °C. The absorbance at 260 nm was determined for each lysate. An equal number of A260 units for each lysate was overlaid on a 10-50% sucrose gradient (10 mM Tris pH 7.26, 60 mM KCl, 10 mM MgCl_2_, 2.5 mM DTT, 0.2 mg/mL heparin, 10 μg/mL cycloheximide). The gradients were subjected to centrifugation at 30,000 RPM for 3 hours at 4 °C. The absorbance at 254 nm was measured along the gradient using a fractionation system (Teledyne ISCO).

### Analysis of rRNA integrity

For analysis of rRNA integrity, total RNA isolated from cells of interest as described above was denatured in 1x RNA Loading Dye (NEB) containing 100 ng/μL ethidium bromide by incubation at 65 °C for 10 minutes. The denatured RNA was resolved on a 1.5% denaturing formaldehyde agarose gel as described previously (Rio, 2015).

### Analysis of CCLE RNAseq Data and ADAR1 Dependency

Raw CCLE RNaseq count data from breast cancer cell lines were normalized by the ‘cpm’ function of ‘edgeR’(Robinson et al., 2010). From the cpm values z-scores were determined for each gene across all cell lines. To determine ‘ISG Core Score’ or ‘ISG Score’ we calculated the median z-score of previously identified ‘Core ISGs’ (Liu et al., 2019) or all ISGs defined by the GSEA/mSigDB hallmark gene set collection (HALLMARK_INTERFERON_ALPHA_RESPONSE and HALLMARK_INTERFERON_GAMMA_RESPONSE) (Liberzon et al., 2015). Molecular subtypes of breast cancer cell lines were defined previously (Marcotte et al., 2016).

### Analysis of TCGA RNAseq Data

Unnormalized RSEM values were normalized by the ‘cpm’ function of edgeR(Robinson et al., 2010). From the cpm values modified z-scores were determined using the following formula.

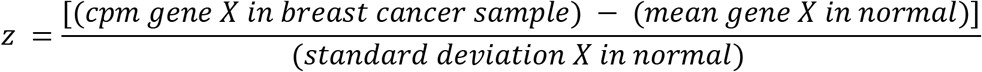

We calculated ‘ISG Core Score’ and ‘ISG Score’ as described above. Molecular subtypes of TCGA samples were defined previously (Lehmann et al., 2016).

### Data and Code Availability

CCLE RNAseq count data (CCLE_RNAseq_genes_counts_20180929.gct.gz, CCLE_RNAseq_rsem_transcripts_tpm_20180929.txt.gz) were obtained from the Broad Institute Cancer Cell Line Encyclopedia and is available online at https://portals.broadinstitute.org/ccle/data. Dependency data (D2_combined_gene_dep_scores.csv, Achilles_gene_effect.csv) were obtained from Broad Institute DepMap Portal and is available on at https://depmap.org/portal/download/. TCGA breast cancer RNAseq (illuminahiseq_rnaseqv2-RSEM_genes, illuminahiseq_rnaseqv2-RSEM_isoforms_normalized) and clinical data (Merge_Clinical) were obtained from the Broad Institute FireBrowse and is available online at http://firebrowse.org/.

All custom R scripts used in this are available upon request. All other data are available in the main text and figures or supplemental information.

### Immunohistochemistry

Human breast formalin fixed paraffin embedded tissue array sections (5μm) on positively charged slides were obtained from US Biomax Inc. (BC081116d). For immunohistochemistry, sections were stained using a Bond RXm autostainer (Leica). Briefly, slides were baked at 65°C for 15min and automated software performed dewaxing, rehydration, antigen retrieval, blocking, primary antibody incubation, post primary antibody incubation, detection (DAB), and counterstaining using Bond reagents (Leica). Samples were then removed from the machine, dehydrated through ethanols and xylenes, mounted and cover-slipped. An antibody for ADAR1-p150 (Abcam ab126745) was diluted 1:100 in Antibody Diluent (Leica).

