## Supplemental Information for "ADAR1 editing dependency in triple-negative breast cancer"

### Supplemental Figure 1

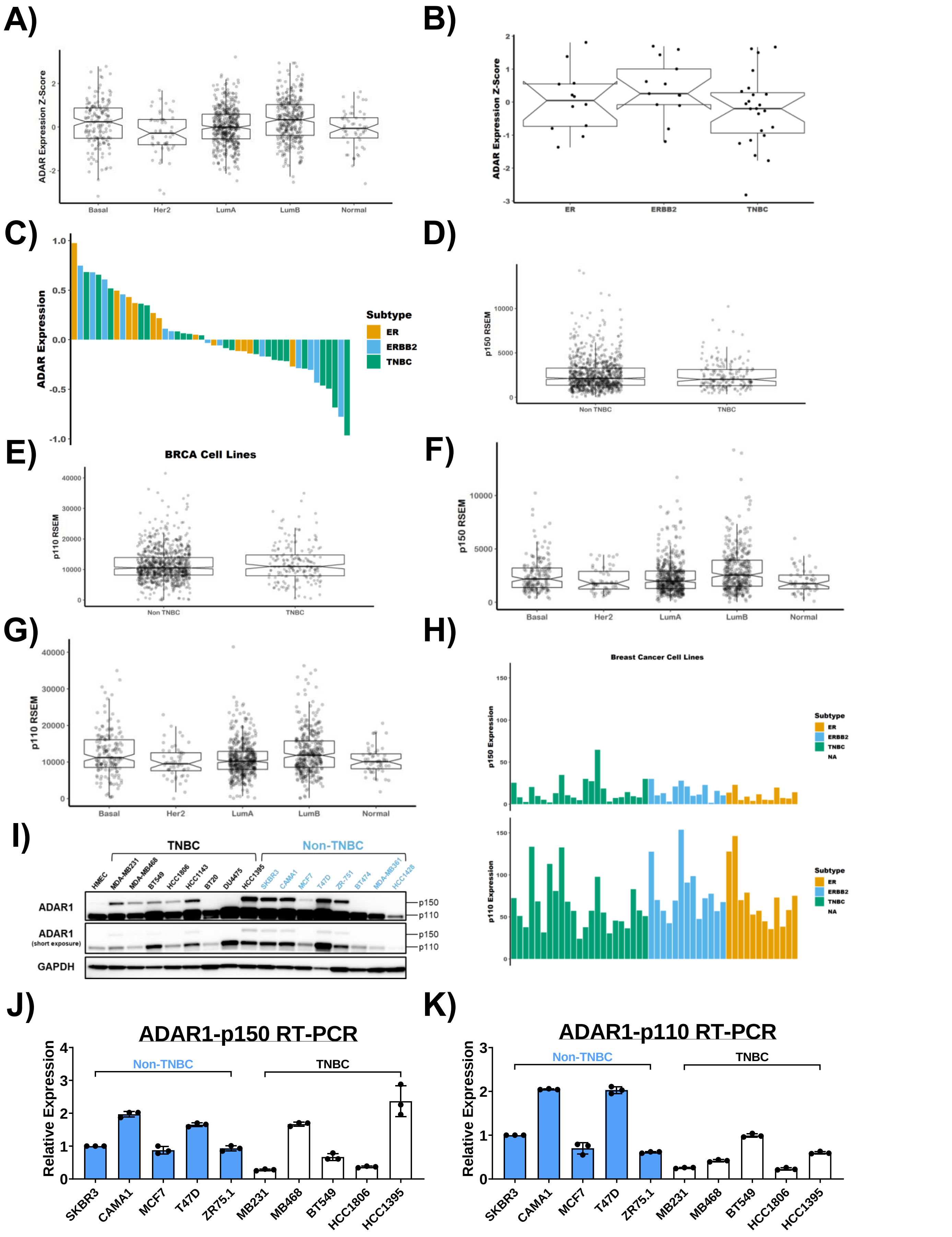

### Supplemental Figure 2

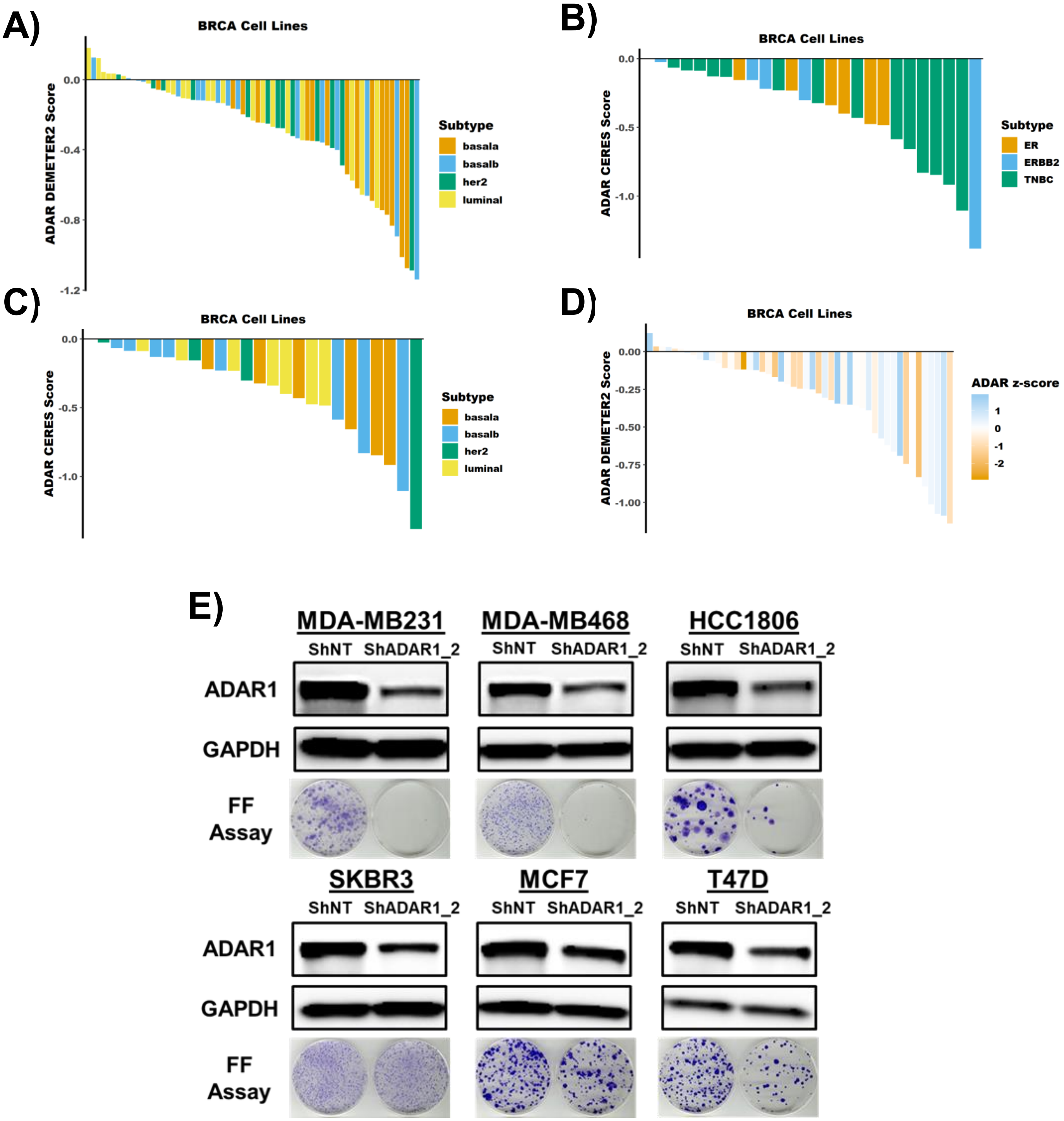

### Supplemental Figure 3

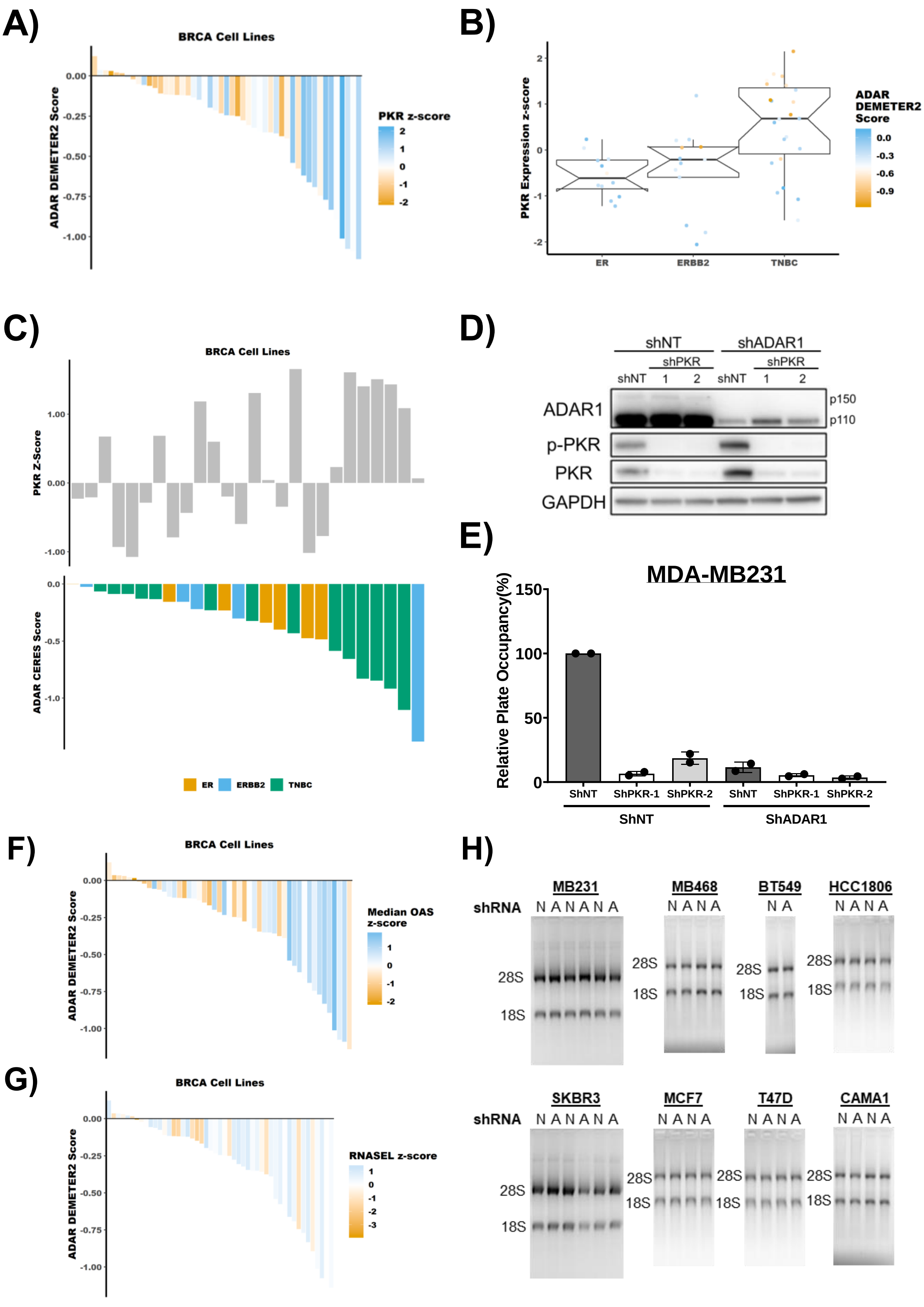

### Supplemental Figure 4

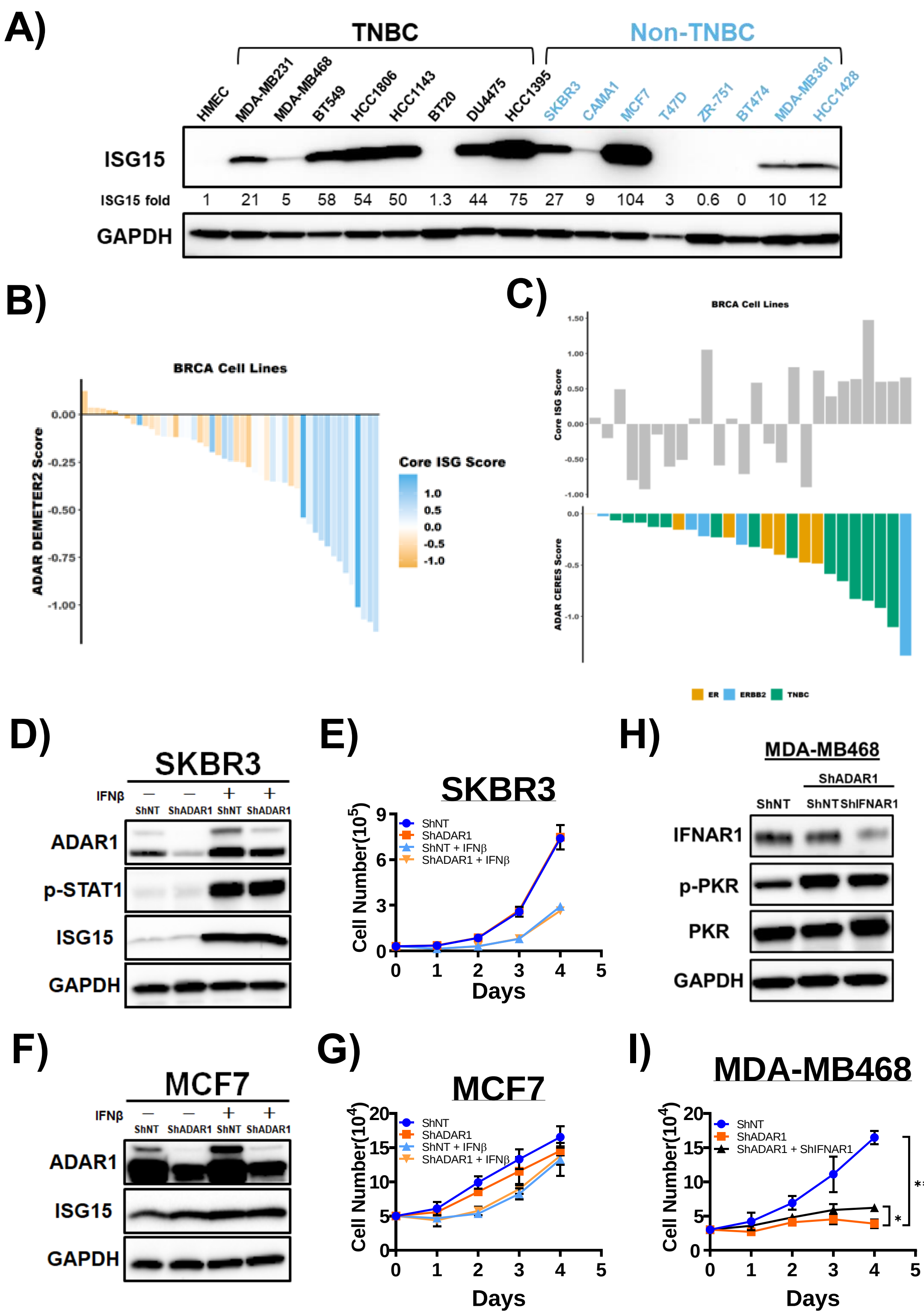

**Supplemental Figure Legends**

**Figure S1. ADAR1 is highly expressed in all breast cancer subtypes. Related to Figure 1.**

A) Relative mRNA expression of ADAR1 in normal, Luminal-A(LumA), Luminal-B(LumB), HER2-positive(Her2) and Basal-like breast cancer. Data were extracted from TCGA database.

B) Relative mRNA expression of ADAR1 in ER-positive, ERBB2(HER2)-positive and TNBC breast cancer cell lines. Data were extracted from CCLE database.

C) Relative protein expression of ADAR1 in breast cancer cell lines. Data were extracted from CCLE database.

D) Relative mRNA expression of ADAR1 p150 isoform in TNBC and Non-TNBC. Data were extracted from TCGA database.

E) Relative mRNA expression of ADAR1 p110 isoform in TNBC and Non-TNBC. Data were extracted from TCGA database.

F) Relative mRNA expression of ADAR1 p150 isoform in normal, Luminal-A(LumA), Luminal-B(LumB), HER2-positive(Her2) and Basal-like breast cancer. Data were extracted from TCGA database.

G) Relative mRNA expression ADAR1 p110 isoform in normal, Luminal-A(LumA), Luminal-B(LumB), HER2-positive(Her2) and Basal-like breast cancer. Data were extracted from TCGA database.

H) Relative mRNA expression of ADAR1 p150 and p110 isoforms in breast cancer cell lines. Data were extracted from CCLE database.

I) Immunoblots showing protein levels of ADAR1 and GAPDH(loading control) in breast cancer cell lines. Images are representative of three replicates.

J) qRT-PCR to detect mRNA levels of ADAR1 p150 isoform in breast cancer cell lines. Gene expression was normalized to GAPDH and compared to SKBR3 set as 1-fold. Data are represented as mean ± SD, N=2.

K) qRT-PCR to detect mRNA levels of ADAR1 p110 isoform in breast cancer cell lines. Gene expression was normalized to GAPDH and compared to SKBR3 set as 1-fold. Data are represented as mean ± SD, N=2.

**Figure S2. ADAR1 is required for TNBC proliferation. Related to Figure 1.**

A) ADAR1-dependency scores in breast cancer cell lines determined by CRISPR-Cas9-mediated ADAR1 reduction. Lower CERES scores indicate stronger ADAR1-dependency.

B) ADAR1-dependency scores in breast cancer cell lines of different subtypes. Lower DEMETER2 scores indicate stronger ADAR1 dependency.

C) ADAR1-dependency scores in breast cancer cell lines of different subtypes, determined by CRISPR-Cas9-mediated ADAR1 reduction. Lower CERES scores indicate stronger ADAR1-dependency.

D) Expression level of ADAR1 doesn’t correlate with ADAR1-dependency scores in breast cancer cell lines. Lower DEMETER2 scores indicate stronger ADAR1-dependency. ADAR1 z-scores indicate ADAR1 expression levels (blue: high; orange: low).

E) Immunoblots showing protein levels of ADAR1 p150 isoform and GAPDH(loading control) with or without ADAR1-knockdown in breast cancer cell lines. Fold change of ADAR1 (ShADAR1_2/ShNT) is indicated, normalized to GAPDH. FF assay showed that ADAR1-knockdown reduced proliferation of TNBC but not Non-TNBC cells. Images are representative, N=2.

**Figure S3. PKR is overexpressed in TNBC and activated upon ADAR loss. Related to Figure 3.**

A) Expression level of PKR correlates with ADAR1-dependency scores in breast cancer cell lines. Lower DEMETER2 scores indicate stronger ADAR1 dependency. PKR z-scores indicate PKR expression levels (blue: high; orange: low).

B) Expression level of PKR correlates with ADAR1-dependency scores in breast cancer cell lines. PKR z-scores indicate PKR expression levels. Lower DEMETER2 scores indicate stronger ADAR1-dependency (blue: low dependency; orange: high dependency).

C) ADAR1-dependency scores positively correlate with PKR expression in breast cancer cell lines. Upper panel: Relative mRNA expression of PKR in breast cancer cell lines. Lower panel: ADAR1-dependency scores determined by CRISPR-Cas9-mediated ADAR1 reduction. Lower CERES scores indicate stronger ADAR1-dependency.

D) Immunoblots showing protein levels of ADAR1, PKR, p-PKR (T446) and GAPDH(loading control) in MDA-MB231 cells. PKR was knocked down in ShADAR1-treated MDA-MB231 cells to determine if PKR loss reverses ADAR1-knockdown phenotype. Images are representative, N=2.

E) Quantification of FF assay with samples from F). Relative plate occupancy was determined using ImageJ software and normalized to ShNT-ShNT. Data are represented as mean ± SD. N=2.

F) Expression level of OAS correlates with ADAR1-dependency scores in breast cancer cell lines. Lower DEMETER2 scores indicate stronger ADAR1 dependency. OAS z-scores indicate OAS expression levels (blue: high; orange: low).

G) Expression level of RNASEL correlates with ADAR1-dependency scores in breast cancer cell lines. Lower DEMETER2 scores indicate stronger ADAR1-dependency. RNASEL z-scores indicate RNASEL expression levels (blue: high; orange: low).

H) Total RNA was extracted from breast cancer cell lines with(A) or without(N) ADAR1-knockdown, and resolved using denaturing agarose gel electrophoresis to detect RNA degradation. The position of 28S and 18S rRNA are noted.

**Figure S4. ADAR1-dependent TNBCs exhibit elevated ISG expression and INFAR1 loss rescues ADAR1 knockdown phenotype. Related to Figure 4.**

A) Immunoblots showing protein levels of ISG15 and GAPDH(loading control) in breast cancer cell lines. Densitometry quantification of gel images was normalized to GAPDH and compared to HMEC signal set as 1-fold. Images are representative from three replicates.

B) ISG core scores correlate with ADAR1-dependency scores in breast cancer cell lines. Lower DEMETER2 scores indicate stronger ADAR1-dependency. ISG core z-scores indicate expression levels of core ISGs (blue: high; orange: low)..

C) ADAR1-dependency scores positively correlate with ISG core scores in breast cancer cell lines. Upper panel: Relative core ISG scores in breast cancer cell lines. Lower panel: ADAR1-dependency scores determined by CRISPR-Cas9-mediated ADAR1 reduction. Lower CERES scores indicate stronger ADAR1-dependency.

D) Immunoblots showing protein levels of ADAR1, p-STAT1 (S727), ISG15 and GAPDH(loading control) in ShNT- or ShADAR1-treated SKBR3 cells 24hr after treatment with vehicle or IFNβ(10ng/ml). Images are representative, N=2.

E) Cell proliferation assay showing that IFNβ treatment did not reduce proliferation of ADAR1-deficient SKBR3 cells over ADAR1-intact cells. Data are represented as mean ± SD. N=2.

F) Immunoblots showing protein levels of ADAR1, ISG15 and GAPDH(loading control) in ShNT- or ShADAR1-treated MCF7 cells 24hr after treatment with vehicle or IFNβ(10ng/ml). Images are representative, N=2.

G) Cell proliferation assay showing that IFNβ treatment did not reduce proliferation of ADAR1-deficient MCF7 cells over ADAR1-intact cells. Data are represented as mean ± SD. N=2.

H) Immunoblots showing protein levels of IFNAR1, PKR, p-PKR (T446) and GAPDH(loading control) in MDA-MB468 cells. IFNAR1 was knocked down in ShADAR1-treated MDA-MB468 cells to determine if IFNAR1 loss reverses ADAR1-knockdown phenotype. Images are representative, N=2.

E) Cell proliferation assay showing that knockdown of IFNAR1 partially rescued ADAR1-knockdown phenotype in MDA-MB468 cells. Data are represented as mean ± SD. N=2. (****) $p<0.0001$. (*) $p<0.05$.

#### Supplemental Table 1. Primers and ShRNAs used in this study

| **Primer for qPCR** | | |
|---|---|---|
| **Target Gene** | **Forward Primer 5'-3'** | **Reverse Primer 5'-3'** |
| *ADAR* (p150) | CAATGCCTCGCGGGCGCAAT | AGCTGTCTGTGCTCATAGCC |
| *ADAR* (p110) | ACTGGCAGTCTCCGGGTG | AGCTGTCTGTGCTCATAGCC |
| *IFNB1* | GCTTCTCCACTACAGCTCTTTC | CAGTATTCAAGCCTCCCATTCA |
| *GAPDH* | GAGTCAACGGATTTGGTCGT | GACAAGCTTCCCGTTCTCAG |
| **Short-hairpin RNA for Lentiviral Transduction** | | |
| **Target Gene** | **ShRNA Sequence** | **Note** |
| *ADAR - 1* | GCCCACTGTTATCTTCACTTT | TRCN0000050788 (MilliporeSigma) |
| *ADAR - 2* | GCTGTTAGAATATGCCCAGTT | TRCN0000050790 (MilliporeSigma) |
| *IFNAR1 - 1* | CCTTAGTGATTCATTCCATATCTC | TRCN0000059014 (MilliporeSigma) |
| *IFNAR1 - 2* | GCCAAGATTCAGGAAATTATTCTC | TRCN0000059013 (MilliporeSigma) |
| *EIF2AK2 - 1* | GAGGCGAGAAACTAGACAAAGCTC | TRCN0000197012 (MilliporeSigma) |
| *EIF2AK2 - 2* | GCTGAACTTCTTCATGTATGTCTC | TRCN0000196400 (MilliporeSigma) |
